# Human NK cell responses to the SARS-CoV-2 Spike_269-277_ peptide YLQPRTFLL

**DOI:** 10.1101/2024.09.26.613632

**Authors:** Eleni Bilev, Simone Schiele, Beatrice Foglietta, Hans-Gustaf Ljunggren, Quirin Hammer

**Affiliations:** Center for Infectious Medicine, Department of Medicine Huddinge, Karolinska Institutet, Karolinska University Hospital Huddinge, Stockholm, Sweden

## Abstract

Natural killer (NK) cells act as the first line of defense against virus infections. The effector functions of human NK cells are controlled by inhibitory and activating receptors, including NKG2A and NKG2C, which recognize peptides presented by HLA-E. Recent studies have suggested that the SARS-CoV-2 Spike_269-277_ peptide YLQPRTFLL may modulate NK cell activity. Here, we show that the YLQPRTFLL peptide is poorly presented by HLA-E. Functional interrogation further revealed that loading of target cells with YLQPRTFLL did not affect the effector functions of NKG2A_+_ nor NKG2C_+_ NK cells. Our findings thus indicate that the Spike_269-277_ peptide YLQPRTFLL has a limited influence on human NK cell responses.

## Introduction

Natural killer (NK) cells are innate lymphoid cells that act as the first line of defense against virus infections. In agreement with their anti-viral functions, NK cells are robustly activated during acute COVID-19 and correlate with clearance of SARS-CoV-2 infection [1-3]. NK cell responses are regulated by numerous receptors, including the inhibitory receptor NKG2A as well as its activating counterpart NKG2C, both of which specifically recognize HLA-E/peptide complexes. Correspondingly, presentation of viral peptides by HLA-E has been demonstrated to regulate NK cell activity [4, 5]. Recent studies have suggested that the SARS-CoV-2 Spike_269-277_ peptide `YLQPRTFLL` may modulate NK cell responses via presentation by HLA-E and subsequent binding of HLA-E/`YLQPRTFLL` complexes to NKG2A or NKG2C [6, 7].

## Results

To explore the impact of the SARS-CoV-2 Spike_269-277_ peptide on NK cells, we first assessed its presentation by HLA-E. For this, we loaded varying concentrations of high-purity synthetic `YLQPRTFLL` peptide on HLA-E*01:03-expressing K562 (K562/HLA-E) cells [8] and monitored the formation of HLA-E/peptide complexes at the cell surface. These experiments revealed that Spike_269-277_ displayed significantly lower presentation by HLA-E compared to the well-known HLA-E-restricted peptides `VMAPRTLIL` (signal peptide of HLA-C) and `VMAPRTLFL` (signal peptide of HLA-G) over a broad range of concentrations (**Figure 1A, B**).

**Figure 1.**
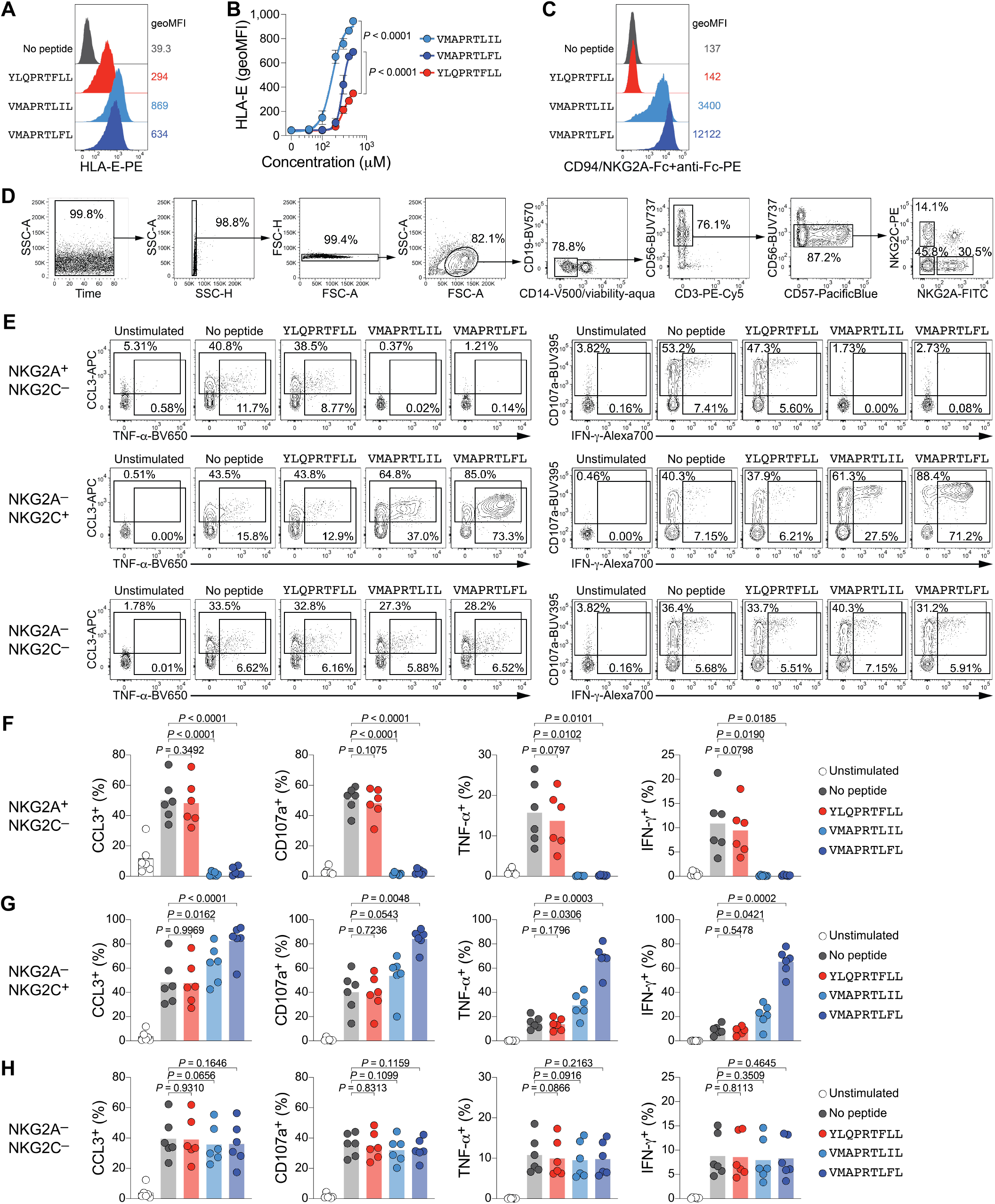
The SARS-CoV-2 Spike_269-277_ peptide YLQPRTFLL does not affect NK cell responses. **(A, B)** K562/HLA-E cells were incubated with solvent control (No peptide) or the indicated peptides over-night and stabilization of HLA-E at the cell surface was detected by flow cytometry. Representative HLA-E staining upon loading with 300 µM peptides. **(B)** Quantification of HLA-E stabilization (pooled data from n=4 independent experiments, displayed as mean±SEM; repeated-measures two-way ANOVA with Dunnett’s multiple comparison test). **(C)** K562/HLA-E were loaded with 500 µM peptides and incubated with recombinant CD94/NKG2A-Fc fusion proteins, followed by flow cytometric detection with anti-Fc antibody (one representative of n=2 independent experiments). **(D-H)** Enriched NK cells from healthy donors were cultured with peptideloaded K562/HLA-E to assess functional responses. **(D)** Gating strategy to identify viable CD14^−^ CD19^−^ CD3^−^ CD56^dim^ NK cell sub-populations with flow cytometry. **(E)** Representative effector functions of NK cell sub-populations cultured alone (Unstimulated) or in the presence of K562/HLA-E treated with solvent control (No peptide) or loaded with 300 µM of the indicated peptides. **(F)** Quantification of effector functions of NKG2A^+^ NKG2C^−^ NK cells, **(G)** NKG2C^+^ NKG2A^−^ NK cells, and **(H)** NKG2A^−^ NKG2C^−^ NK cells (n=6 donors in n=4 independent experiments, circles indicate individual donors and bars represent mean, repeated-measures one-way ANOVA with Dunnett’s multiple comparisons test).

Next, we loaded K562/HLA-E cells with saturating amounts of peptide and evaluated the binding of recombinant CD94/NKG2A receptors to the peptide-loaded cells. While presentation of the control peptides `VMAPRTLIL`, and especially `VMAPRTLFL`, resulted in clear receptor binding, loading with `YLQPRTFLL` did not lead to detectable binding of CD94/NKG2A (**Figure 1C**).

To directly interrogate functional responses of human NK cells against Spike_269-277_, we cultured enriched NK cells from healthy donors with peptide-loaded K562/HLA-E cells and determined degranulation as well as chemokine and cytokine expression (**Figure 1D, E**). In these assays, CD56^dim^ NK cells expressing the inhibitory receptor NKG2A and lacking NKG2C were fully inhibited by the two control peptides `VMAPRTLIL` and `VMAPRTLFL` (**Figure 1F**). In line with previous data, the effector functions of NKG2A^−^ NKG2C^+^ CD56^dim^ NK cells were robustly triggered in response to `VMAPRTLIL` and particularly `VMAPRTLFL`-loaded target cells (**Figure 1G**) [4]. As expected, NKG2A^−^ NKG2C^−^ CD56^dim^ NK cells that do not express HLA-E/peptide-specific receptors did not show differential activity (**Figure 1H**). Importantly, loading of K562/HLA-E with the SARS-CoV-2 Spike_269-277_ peptide `YLQPRTFLL` did not have a significant impact on the activity of either NKG2A^+^ NKG2C^−^ or NKG2A^−^ NKG2C^+^ NK cells (**Figure 1F, G**).

Taken together, we find poor presentation of Spike_269-277_ by HLA-E, no binding to recombinant NK cell receptors, and no impact on the effector functions of NKG2A^+^ nor NKG2C^+^ NK cells *in vitro*. Thus, these data indicate that the Spike_269-277_ peptide `YLQPRTFLL` has a limited influence on NK cell responses.

## Discussion

In this study, we find poor presentation of the SARS-CoV-2 Spike_269-277_ peptide `YLQPRTFLL` by HLA-E, no binding of HLA-E/`YLQPRTFLL` complexes to recombinant NK cell receptors, and no impact on the *in vitro* effector functions of NK cells, which is in contrast to recently reported observations [6, 7]. What could cause the discrepancies between our findings and those of previous studies? While the exact reasons are currently unknown, it cannot be excluded that experimental differences may account for these distinct observations.

Hasan *et al*. used the anti-HLA-E antibody clone 4D12 to investigate HLA-E expression [7]. Clone 4D12 was previously shown to preferentially bind peptide-free forms of HLA-E, the levels of which decrease upon peptide loading [9]. In contrast, clone 3D12, which was used in the present study, detects HLA-E/peptide complexes and its binding increases upon peptide loading [9], thus accurately reporting the presentation of HLA-E-restricted peptides. Furthermore, the finding that the SARS-CoV-2 Spike_269-277_ peptide `YLQRPTFLL` displays a low capacity to stabilize HLA-E on the surface of K562/HLA-E cells, compared to the two well-characterized positive control peptides `VMAPRTLIL` and `VMAPRTLFL`, is in agreement with results from McMichael and colleagues, who found comparably low binding of `YLQRPTFLL` to HLA-E in UV-exchange ELISA, when expressed as single chain trimer, and in thermal stability assays [10].

K562/HLA-E cells are a sensitive tool for peptide stabilization assays since they do not express other HLA class I molecules [8]. In contrast, T2 cells express HLA-A*02 [11], which may interfere with peptide presentation because certain peptides can be bound by both HLA-E and HLA-A*02 [12]. HLA-A*02-restricted T cell responses against the Spike_269-277_ peptide `YLQPRTFLL` have indeed been reported after SARS-CoV-2 infection as well as BNT162b2 vaccination [13, 14]. Given these considerations, the use of enriched NK cells as effectors in short-term assays may be critical to exclude unintended peptide-driven activation of HLA-A*02-restricted T cells, which could indirectly promote NK cell functions via release of soluble factors or activation of accessory cells. Additionally, the absence of other HLA class I molecules such as HLA-B or HLA-C on K562/HLA-E cells prevents their recognition by inhibitory NK cell receptors other than NKG2A, thereby allowing for specifically determining the functional impact of the loaded peptides. Instead, co-culture of NK cells with target cells that express classical HLA class I molecules, such as Calu-3, has the potential to generate either matched or mis-matched settings.

For instance, NKG2A^+^ NK cells that co-express KIR2DL1 would not only detect differences in HLA-E-presented peptides but could also be inhibited by HLA-C*15:02 on Calu-3 cells [15]. Correspondingly, if NKG2C^+^ NK cells co-express KIR2DL3 while the cognate HLA-C ligands are absent on Calu-3 [15], this would be expected to result in missing-self recognition, possibly enhancing responses independently of HLA-E. Such complexities likely further increase upon infection with SARS-CoV-2 and require additional stringent controls to avoid HLA-E-independent contributions due to infection-induced alterations in the expression of classical HLA class I molecules.

The use of synthetic peptides for loading of K562/HLA-E cells could be considered a limitation of our study since it does not provide insights into the processing cascade upstream of presentation by HLA-E. Bortolotti *et al*. transfected Spike protein into K562 and Beas-B2 cells, reported enhanced HLA-E expression on the treated cells, and predicted that this could be driven by the octamer `LQPRTFLL` [6], part of the Spike_269-277_ peptide `YLQPRTFLL`. Hasan *et al*. used transfection of Calu-3 and 721.221 cells for transient delivery of plasmid DNA encoding full-length Spike [7]. These experimental settings incorporate multiple steps prior to peptide presentation. Conversely, loading of K562/HLA-E cells with exogenous candidate peptides enables the sensitive assessment of presentation over a broad range of defined concentrations, as it bypasses variability in transfection efficiency, DNA construct design, antigen expression, and potential rate-limiting steps in proteasomal degradation or translocation into the endoplasmic reticulum by transporter associated with antigen processing [16].

Collectively, by using an antibody specific for HLA-E/peptide complexes and functional assays that are not influenced by non-NK immune cells or other HLA molecules, we found no striking evidence that the SARS-CoV-2 Spike_269-277_ peptide `YLQPRTFLL` controls human NK cell responses *in vitro*. Thus, these findings suggest that further studies are required to clarify whether Spike-derived peptides influence the activity of NK cell sub-populations against SARS-CoV-2.

## Acknowledgements

We thank Timo Rückert and Josefine Dunst for critical feedback. We are grateful to all members of the Center for Infectious Medicine for inspiring discussions. This work was supported by Åke Wibergs Stiftelse (M22-0013), KI Foundations (2022-01606), KI Foundation for Virus Research (2021-00069, 2022-00245, and 2023-00155), Petrus och Augusta Hedlunds Stiftelse (M2021-1533 and M2022-1821), Stiftelsen Clas Groschinskys Minnesfold (M21120 and M2233), Stiftelsen Lars Hiertas Minne (FO2021-0263 and FO2023-0167), Stiftelsen Tornspiran, and Jonas Söderquist Stiftelse (all to Q.H.). This manuscript was typeset with the bioRxiv word template by @Chrelli: www.github.com/chrelli/bioRxiv-word-template

## Author contributions

E.B., S.S., and B.F. performed experiments, and analyzed as well as visualized the data. H.-G.L. provided resources. Q.H. conceptualized the study, acquired funding, supervised the work, and administered the project. E.B., H.-G.L., and Q.H. wrote the original draft and all authors reviewed, edited and approved the manuscript.

## Materials and Methods

### Cells and cell lines

K562 cells expressing HLA-E_∗_01:03 (K562/HLA-E cells; generated by E. Weiss, Ludwig Maximilian University [8]) were maintained in complete RPMI (RPMI-1640 supplemented with 2 mM glutamine, 10% [v/v] FBS, 100 U/mL penicillin, and 100 μg/mL streptomycin; all Gibco) supplemented with 1 mg/mL G418 (Gibco).

Buffy coats from healthy donors were obtained from the Department of Clinical Immunology and Transfusion Medicine, Karolinska University Hospital as approved by the Ethical Review Board Stockholm (DNR 2020-05289). PBMC were isolated from buffy coats with standard density gradient centrifugation, cryopreserved in FBS containing 10% (v/v) DMSO, and thawed for magnetic enrichment of NK cells using NK Cell Isolation Kit (Miltenyi Biotec) as described previously [17].

### Flow cytometry

Antibody staining and flow cytometric analyses were performed following established guidelines [18]. In brief, cell suspensions were incubated with combinations of fluorochrome-conjugated antibodies (**Table S1**) at optimized concentrations in PBS for 20 min at RT. Dead cells were identified with LIVE/DEAD Fixable Aqua Dead Cell Stain Kit (ThermoFisher) and excluded from analyses. All samples were fixed before acquisition using BD Cytofix/Cytoperm (BD Biosciences) according to the manufacturer’s instructions and, if indicated, intracellular proteins were stained with fluorochrome-conjugated antibodies in Perm/Wash buffer (BD Biosciences) for 30 min at 4°C. Samples were acquired on an LSR Fortessa flow cytometer (BD Biosciences) and analyzed with FlowJo v10.10.0 (BD Biosciences).

### Peptide loading

K562/HLA-E cells were loaded with peptides as described previously [4, 5]. In brief, synthetic custom peptides of ≥95% purity (Peptides&Elephants) were reconstituted in sterile water at a stock concentration of 12 mM and K562/HLA-E cells were seeded at 2×106 cells/mL in serum-free Opti-MEM (ThermoFisher). If not indicated otherwise, synthetic peptides were added at a final concentration of 300 μM to the cells and incubated at 37°C overnight (15-18 h). The corresponding volume of sterile water was added to separate wells as solvent control (referred to as no peptide). After over-night loading, peptide-loaded cells were either washed with complete RPMI and used in co-cultures with NK cells for functional assays (see below) or washed with PBS and stained for flow cytometric analysis of HLA-E using antibody clone 3D12 (**Table S1**). For assessing receptor binding, K562/HLA-E cells loaded with 500 μM peptides were incubated with 50 μg/mL recombinant CD94/NKG2A-Fc fusion protein (Acro Biosystems) for 45 min on ice, followed by secondary staining with anti-human Fc antibody (**Table S1**).

### Functional assays

To investigate functional responses, magnetically enriched NK cells were rested in complete RPMI at 1×10^6^ cells/mL over-night (15-18 h) and on the next day, 1×10^5^ NK cells were co-cultured with 1×10^5^ peptide-loaded K562/HLA-E for 6 h in 200 μL complete RPMI in V-bottom 96-well plates at 37°C. Peptides were added to the co-cultures at a final concentration of 300 μM. To detect degranulation, anti-CD107a antibody (**Table S1**) was added at the start of the co-culture and GolgiStop as well as GolgiPlug (containing monensin and brefeldin A, respectively, both BD Biosciences) were added after 1 h. After 6 h total co-culture time, the cells were stained for surface markers (**Table S1**) and fixed (BD Cytofix/Cytoperm, BD Biosciences). Fixed cells were permeabilized for intracellular detection of CCL3, IFN-`γ`, and TNF-α (**Table S1**) in Perm/Wash buffer (BD Biosciences). NK cells were identified as single viable CD14^−^ CD19^−^ CD3^−^ CD56^dim^ cells and further stratified for sub-populations based on the expression of NKG2A and NKG2C, as displayed in the gating strategy in **Figure 1E**.

**Supplementary Table S1.** Antibodies used in this study.

| Antibody (clone) | Manufacturer | RRID |
| --- | --- | --- |
| Anti-CCL3-APC (REA257) | Miltenyi Biotec | AB_2651376 |
| Anti-CD107a-BUV395 (H4A3) | BD Biosciences | AB_2739073 |
| Anti-CD14-V500 (M5E2) | BD Biosciences | AB_10611856 |
| Anti-CD19-BV570 (HIB19) | BioLegend | AB_2563606 |
| Anti-CD3-PE-Cy5 (UCHT1) | Beckman Coulter | AB_2827419 |
| Anti-CD56-BUV737 (NCAM16.2) | BD Biosciences | AB_2860005 |
| Anti-CD16-BV785 (3G8) | BioLegend | AB_2563803 |
| Anti-CD57-PacificBlue (HNK-1) | BioLegend | AB_2562459 |
| Anti-HLA-E-PE (3D12) | BioLegend | AB_1659249 |
| Anti-IFN-γ-Alexa700 (B27) | BD Biosciences | AB_396977 |
| Anti-NKG2A-ViobrightFITC (REA110) | Miltenyi Biotec | AB_2726173 |
| Anti-NKG2C-PE (REA205) | Miltenyi Biotec | AB_2751835 |
| Anti-TNF-α-BV650 (MAb11) | BioLegend | AB_2562741 |

